# Detecting high-order epistasis in nonlinear genotype-phenotype maps

**DOI:** 10.1101/072256

**Authors:** Zachary R. Sailer, Michael J. Harms

## Abstract

High-order epistasis has been observed in many genotype-phenotype maps. These multi-way interactions between mutations may be useful for dissecting complex traits and could have profound implications for evolution. Alternatively, they could be a statistical artifact. High-order epistasis models assume the effects of mutations should add, when they could in fact multiply or combine in some other nonlinear way. A mismatch in the “scale” of the epistasis model and the scale of the underlying map would lead to spurious epistasis. In this paper, we develop an approach to estimate the nonlinear scales of arbitrary genotype-phenotype maps. We can then linearize these maps and extract high-order epistasis. We investigated seven experimental genotype-phenotype maps for which high-order epistasis had been reported previously. We find that five of the seven maps exhibited nonlinear scales. Interestingly, even after accounting for nonlinearity, we found statistically significant high-order epistasis in all seven maps. The contributions of high-order epistasis to the total variation ranged from 2.2% to 31.0%, with an average across maps of 12.7%. Our results provide strong evidence for extensive high-order epistasis, even after nonlinear scale is taken into account. Further, we describe a simple method to estimate and account for nonlinearity in genotype-phenotype maps.

## Introduction

Recent analyses of genotype-phenotype maps have revealed “high-order” epistasis—that is, interactions between three, four, and even more mutations (Ritchie et al. 2001; Segrè et al. 2005; Xu et al. 2005; Tsai et al. 2007; Imielinski and Belta 2008; Matsuura et al. 2009; da Silva et al. 2010; Pettersson et al. 2011; Wang et al. 2012; Weinreich et al. 2013; Hu et al. 2013; Sun et al. 2014; Anderson et al. 2015; Yokoyama et al. 2015). The importance of these interactions for understanding biological systems and their evolution is the subject of current debate (Poelwijk et al. 2016; Weinreich et al. 2013). Can they be interpreted as specific, biological interactions between loci? Or are they misleading statistical correlations?

We set out to tackle one potential source of spurious epistasis: a mismatch between the “scale” of the map and the scale of the model used to dissect epistasis (Fisher 1918; Rothman et al. 1980; Frankel and Schork 1996; Cordell 2002; Phillips 2008; Szendro et al. 2013). The scale defines how to combine mutational effects. On a linear scale, the effects of individual mutations are added. On a multiplicative scale, the effects of mutations are multiplied. Other, arbitrarily complex scales, are also possible (Rokyta et al. 2011; Schenk et al. 2013; Blanquart 2014).

Application of a linear model to a nonlinear map will lead to apparent epistasis (Fisher 1918; Rothman et al. 1980; Frankel and Schork 1996; Cordell 2002; Phillips 2008; Szendro et al. 2013). Consider a map with independent, multiplicative mutations. Analysis with a multiplicative model will give no epistasis. In contrast, analysis with a linear model will give epistatic coefficients to account for the multiplicative nonlinearity (Cordell 2002; Phillips 2008). Epistasis arising from a mismatch in scale is mathematically valid, but obscures a key feature of the map: its scale. It is also not parsimonious, as it uses many coefficients to describe a potentially simple nonlinear function. Finally, it can be misleading because these epistatic coefficients partition global information about the nonlinear scale into (apparently) specific interactions between mutations.

Most high-order epistasis models assume a linear scale (or a multiplicative scale transformed onto a linear scale) (Heckendorn and Whitley 1999; Szendro et al. 2013; Weinreich et al. 2013; Poelwijk et al. 2016). These models sum the independent effects of mutations to predict multi-mutation phenotypes. Epistatic coefficients account for the difference between the observed phenotypes and the phenotypes predicted by summing mutational effects. The epistatic coefficients that result are, by construction, on the same linear scale (Poelwijk et al. 2016; Weinreich et al. 2013; Heckendorn and Whitley 1999).

Because the underlying scale of genotype-phenotype maps is not known a priori, the interpretation of high-order epistasis extracted on a linear scale is unclear. If a nonlinear scale can be found that removes high-order epistasis, it would suggest that high-order epistasis is spurious: a highly complex description of a simple, nonlinear system. In contrast, if no such scale can be found, high-order epistasis provides a window into the profound complexity of genotype-phenotype maps.

In this paper, we set out to estimate the nonlinear scales of experimental genotype-phenotype maps. We then account for these scales in the analysis of high-order epistasis. We took our inspiration from the treatment of multiplicative maps, which can be transformed into linear maps using a log transform. Along these same lines, we set out to transform genotype-phenotype maps with arbitrary, nonlinear scales onto a linear scale for analysis of high-order epistasis. We develop our methodology using simulations and then apply it to experimentally measured genotype-phenotype maps.

## Materials and Methods

### Experimental data sets

We collected a set of published genotype-phenotype maps for which high-order epistasis had been reported previously. Measuring an L^th^-order interaction requires knowing the phenotypes of all binary combinations of L mutations—that is, 2^L^ genotypes. The data sets we used had exhaustively covered all 2^L^ genotypes for five or six mutations. These data sets cover a broad spectrum of genotypes and phenotypes. Genotypes included point mutations to a single protein (Weinreich et al. 2006), point mutations in both members of a protein/DNA complex (Anderson et al. 2015), random genomic mutations (Khan et al. 2011; de Visser et al. 2009), and binary combinations of alleles within a biosynthetic network (Hall et al. 2010). Measured phenotypes included selection coefficients (Weinreich et al. 2006; Khan et al. 2011; de Visser et al. 2009), molecular binding affinity (Anderson et al. 2015), and yeast growth rate (Hall et al. 2010). (For several data sets, the “phenotype” is a selection coefficient. We do not differentiate fitness from other properties for our analyses; therefore, for simplicity, we will refer to all maps as genotype-phenotype maps rather than specifying some as genotype-fitness maps). All data sets had a minimum of three independent measurements of the phenotype for each genotype. All data sets are available in a standardized ascii text format.

### Nonlinear scale

We described nonlinearity in the genotype-phenotype map by a power transformation (see Results) (Box and Cox 1964; Carroll and Ruppert 1981). The independent variable for the transformation was 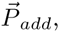, the predicted phenotypes of all genotypes assuming linear and additive affects for each mutation. The estimated additive phenotype of genotype *i*, is given by:

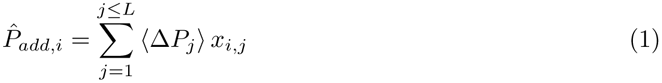

where 〈∆*P*_*j*_〉 is the average effect of mutation *j* across all backgrounds, *x*_*i, j*_ is an index that encodes whether or not mutation *j* is present in genotype *i*, and *L* is the number of sites. The dependent variables are the observed phenotypes 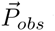 taken from the experimental genotype-phenotype maps. We use nonlinear least-squares regression to fit and estimate the power transformation from 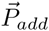 to 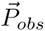

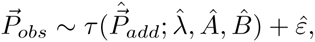

where ε is a residual and τ is a power transform function. This is given by:

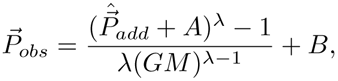

where *A* and *B* are translation constants, *GM* is the geometric mean of 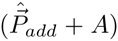, and λ is a scaling parameter. We used standard nonlinear regression techniques to minimize *d:*

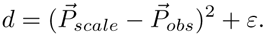

We then reversed this transformation to linearize *P*_*obs*_ using the estimated parameters 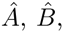 and 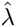. We did so by the back-transform:

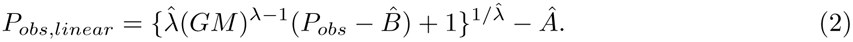

### High-order epistasis model

We dissected epistasis using a linear, high-order epistasis model. These have been discussed extensively elsewhere (Heckendorn and Whitley 1999; Poelwijk et al. 2016; Weinreich et al. 2013), so we will only briefly and informally review them here.

A high-order epistasis model is a linear decomposition of a genotype-phenotype map. It yields a set of coefficients that account for all variation in phenotype. The signs and magnitudes of the epistatic coefficients quantify the effect of mutations and interactions between them. A binary map with 2^*L*^ genotypes requires 2^*L*^ epistatic coefficients and captures all interactions, up to *L^th^-order*, between them. This is conveniently described in matrix notation.

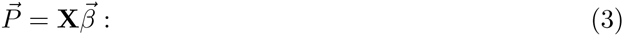

a vector of phenotypes 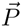 can be transformed into a vector of epistatic coefficients 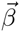 using a 2^*L*^ × 2^*L*^ decomposition matrix that encodes which coefficients contribute to which phenotypes. If X is invertible, one can determine 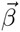 from a collection of measured phenotypes by

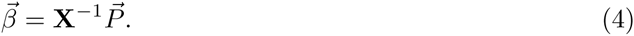

**X** can be formulated in a variety of ways (Poelwijk et al. 2016). Following others in the genetics literature, we use the form derived from Walsh polynomials (Heckendorn and Whitley 1999; Weinreich et al. 2013; Poelwijk et al. 2016). In this form, **X** is a Hadamard matrix. Conceptually, the transformation identifies the geometric center of the genotype-phenotype map and then measures the average effects of each mutation and combination of mutations in this “average” genetic background (Fig 1). To achieve this, we encoded each mutation at each site in each genotype as -1 (wildtype) or +1 (mutant) (Heckendorn and Whitley 1999; Weinreich et al. 2013; Poelwijk et al. 2016). This has been called a Fourier analysis,(Szendro et al. 2013; Neidhart et al. 2013), global epistasis (Poelwijk et al. 2016), or a Walsh space (Heckendorn and Whitley 1999; Weinreich et al. 2006). Another common approach is to use a single wildtype genotype as a reference and encode mutations as either 0 (wildtype) or 1 (mutant) (Poelwijk et al. 2016).

**Fig 1:**
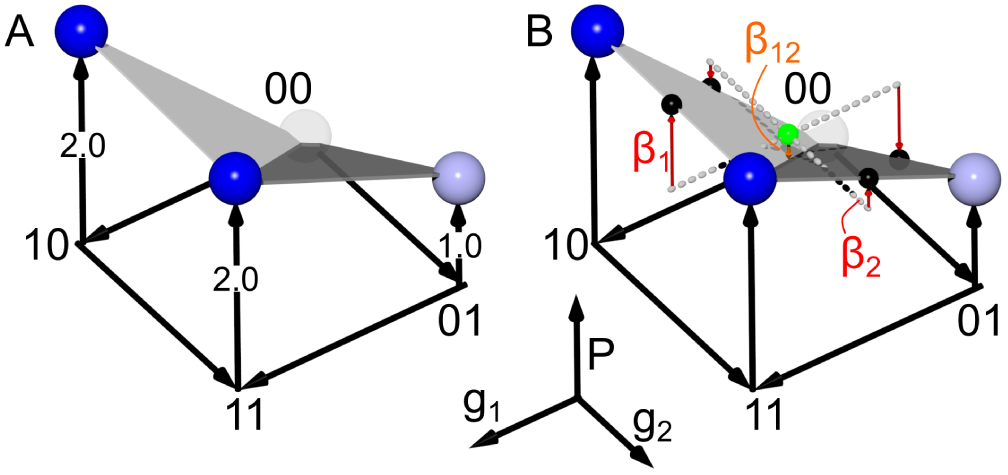
Epistasis can be quantified using Walsh polynomials. A) A genotype-phenotype map exhibiting negative epistasis. Axes are genotype at position 1 (*g*_1_), genotype at position 2 (*g*_2_), and phenotype (*P*). For genotypic axes, “0” denotes wildtype and “1” denotes a mutant. Phenotype is encoded both on the P-axis and as a spectrum from white to blue. The map exhibits negative epistasis: relative to wildtype, the effect of the mutations together (*P*_11_ = 2) is less than the sum of the individual effects of mutations (*P*_10_ + *P*_01_ = 1 + 2 = 3). B) The map can be decomposed into epistatic coefficients using a Walsh polynomial, which measures the effects of each mutation relative to the geometric center of the genotype-phenotype map (green sphere). The additive coefficients *β*_1_ and *β*_2_ (red arrows) are the average effect of each mutation in all backgrounds. The epistatic coefficient *β*_1_2 (orange arrow) the variation not accounted for by *β*_1_ and *β*_2_. Geometrically, it is the distance between the center of the map and the “fold” given by vector connecting *P*_00_ and *P*_11_.

One data set (IV, Table I) has four possible states (A, G, C and T) at two of the sites. We encoded these using the WYK tetrahedral-encoding scheme(Zhang and Zhang 1991; Anderson et al. 2015). Each state is encoded by a three-bit state. The wildtype state is given the bits (1,1,1). The remaining states are encoded with bits that form corners of a tetrahedron. For example, the wildtype of site 1 is G and encoded as the (1,1,1) state. The remaining states are encoded as follows: A is (1, −1, −1), C is (−1, 1, −1) and T is (1, −1, −1).

**Table 1.**

| **Table 1** |  |  |  |  |
| --- | --- | --- | --- | --- |
| ID | genotype | phenotype | $L$ | reference |
| I | scattered genomic mutations | *E. coli* fitness | 5 | (Khan et al. 2011) |
| II | chromosomes in asexual fungi | *A. niger* fitness | 5 | (de Visser et al. 2009) |
| III | protein point mutants | bacterial fitness | 5 | (Weinreich et al. 2006) |
| IV | DNA/protein point mutants | *in vitro* DNA/protein binding affinity | 5 | (Anderson et al. 2015) |
| V | chromosomes in asexual fungi | *A. niger* fitness | 5 | (de Visser et al. 2009) |
| VI | alleles in biosynthetic network | *S. cerevisiae* haploid growth rate | 6 | (Hall et al. 2010) |
| VII | alleles in biosynthetic network | *S. cerevisiae* diploid growth rate | 6 | (Hall et al. 2010) |

### Experimental uncertainty

We used a bootstrap approach to propagate uncertainty in measured phenotypes into uncertainty in epistatic coefficients. To do so we: 1) calculated the mean and standard deviation for each phenotype from the published experimental replicates; 2) sampled the uncertainty distribution for each phenotype to generate a pseudoreplicate vector 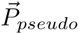 that had one phenotype per genotype; 3) rescaled 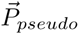 using a power-transform; and 4) determined the epistatic coefficients for 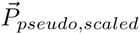 We then repeated steps 2-4 until convergence. We determined the mean and variance of each epistatic coefficient after every 50 pseudoreplicates. We defined convergence as the mean and variance of every epistatic coefficient changed by < 0.1 % after addition of 50 more pseduoreplicates. On average, convergence required ≈ 100, 000 replicates per genotype-phenotype map. Finally, we used a z-score to determine if each epistatic coefficient was significantly different than zero. To account for multiple testing, we applied a Bonferroni correction to all p-values (Abdi 2007).

### Computational methods

Our full epistasis software package—written in Python3 extended with Numpy and Scipy (van der Walt et al. 2011)—is available for download via github (https://harmslab.github.com/epistasis). We used the python package scikit-learn for all regression (Pedregosa et al. 2011). Plots were generated using matplotlib and jupyter notebooks (Hunter 2007; Perez and Granger 2007).

## Results & Discussion

### Nonlinear scale induces apparent high-order epistasis

Our first goal was to understand how a nonlinear scale, if present, would affect estimates of high-order epistasis. To probe this question, we constructed a five-site binary genotype-phenotype map on a nonlinear scale, and then extracted epistasis assuming a linear scale. The nonlinear scale we chose was saturating function:

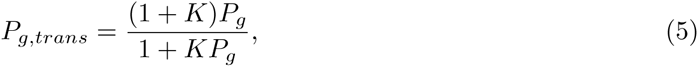

where *P*_*g*_ is the linear phenotype of genotype *g, P*_*g, trans*_ is the transformed phenotype of genotype *g*, and *K* is a scaling constant. As *K* → 0, the map becomes linear. As *K* increases, mutations have systematically smaller effects when introduced into backgrounds with higher phenotypes.

We calculated *P*_*g*_ for all 2^*L*^ binary genotypes using the random, additive coefficients shown in Fig 2A. These coefficients included no epistasis. We then transformed *P*_*g*_ onto the nonlinear *P*_*g, trans*_ scale using Equation 5 with the relatively shallow (*K* = 2) saturation curve shown in Fig 2B. Finally, we applied a linear epistasis model to *P*_*g, trans*_ to extract epistatic coefficients.

**Fig 2:**
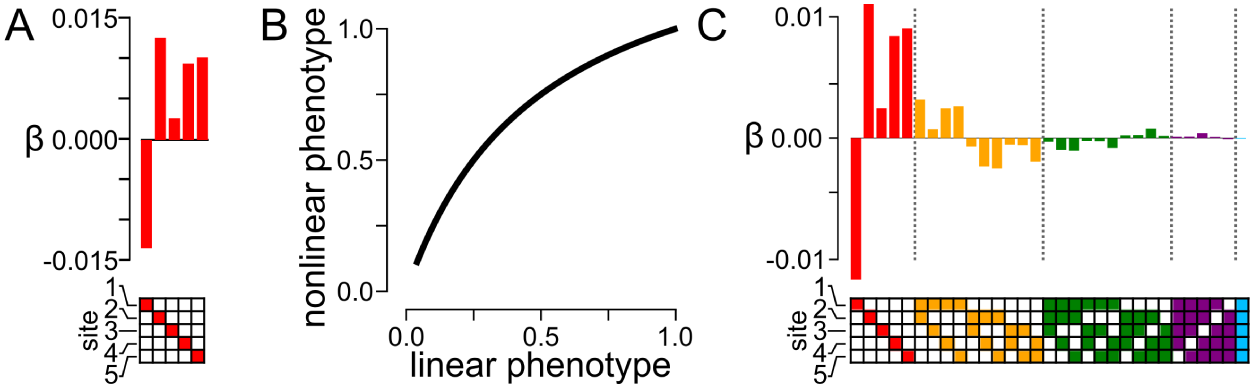
Nonlinearity in phenotype creates spurious high-order epistatic coefficients. A) Simulated, random, first-order epistatic coefficients. The mutated site is indicated by panel below the bar graph; bar indicates magnitude and sign of the additive coefficient. B) A nonlinear map between a linear phenotype and a saturating, nonlinear phenotype. The first-order coefficients in panel A are used to generate a linear phenotype, which is then transformed by the function shown in B. C) Epistatic coefficients extracted from the genotype-phenotype map generated in panels A and B. Bars denote coefficient magnitude and sign. Color denotes the order of the coefficient: first (*β*_*i*_, red), second (*β*_*ij*_, orange), third (*β*_*ijk*_, green), fourth (*β*_*ijkl*_, purple), and fifth (*β*_*ijklm*_, blue). Filled squares in the grid below the bars indicate the identity of mutations that contribute to the coefficient.

We found that nonlinearity in the genotype-phenotype map induced extensive high-order epistasis when the nonlinearity was ignored (Fig 2C). We observed epistasis up to the fourth order, despite building the map with purely additive coefficients. This result is unsurprising: the only mechanism by which a linear model can account for variation in phenotype is through epistatic coefficients (Rothman et al. 1980; Frankel and Schork 1996; Cordell 2002). When given a nonlinear map, it partitions the variation arising from nonlinearity into specific interactions between mutations. This high-order epistasis is mathematically valid, but does not capture the major feature of the map—namely, saturation. Indeed, this epistasis is deceptive, as it is naturally interpreted as specific interactions between mutations. For example, this analysis identifies a specific interaction between mutations one, two, four, and five (Fig 2C, purple). But this four-way interaction is an artifact of the nonlinearity in phenotype of the map, rather than a specific interaction.

### Nonlinear scale and specific epistatic interactions induce different patterns of non-additivity

Our next question was whether we could separate the effects of nonlinear scale and high-order epistasis in binary maps. One useful approach to develop intuition about epistasis is to plot the the observed phenotypes (*P*_*obs*_) against the predicted phenotype of each genotype, assuming linear and additive mutational effects (*P*_*add*_) (Rokyta et al. 2011; Schenk et al. 2013; Szendro et al. 2013). In a linear map without epistasis, *P*_*obs*_ equals *P*_*add*_, because each mutation would have the same, additive effect in all backgrounds. If epistasis is present, phenotypes will diverge from the *P*_*obs*_ = *P*_*add*_ line.

We simulated maps including varying amounts of linear, high-order epistasis, placed them onto increasingly nonlinear scales, and then constructed *P*_*obs*_ vs. *P*_*add*_ plots. We added high-order epistasis by generating random epistatic coefficients and then calculating phenotypes using Eq. 3. We introduced nonlinearity by transforming these phenotypes with Eq. 5. For each genotype in these simulations, we calculated *P*_*add*_ as the sum of the first-order coefficients used in the generating model. *P*_*obs*_ is the observable phenotype, including both high-order epistasis and nonlinear scale.

High-order epistasis and nonlinear scale had qualitatively different effects on *P*_*obs*_ vs. *P*_*add*_ plots. Fig 3A shows plots of *P*_*obs*_ vs. *P*_*add*_ for increasing nonlinearity (left-to-right) and high-order epistasis (bottom-to-top). As nonlinearity increases, *P*_*obs*_ curves systematically relative to *P*_*add*_. This reflects the fact that *P*_*add*_ is on a linear scale and *P*_*obs*_ is on a saturating, nonlinear scale. The shape of the curve reflects the map between the linear and saturating scale: the smallest phenotypes are underestimated and the largest phenotypes overestimated. In contrast, high-order epistasis induces random scatter away from the *P*_*obs*_ = *P*_*add*_ line. This is because the epistatic coefficients used to generate the map are specific to each genotype, moving observations off the expected line, even if the scaling relationship is taken into account.

**Fig 3:**
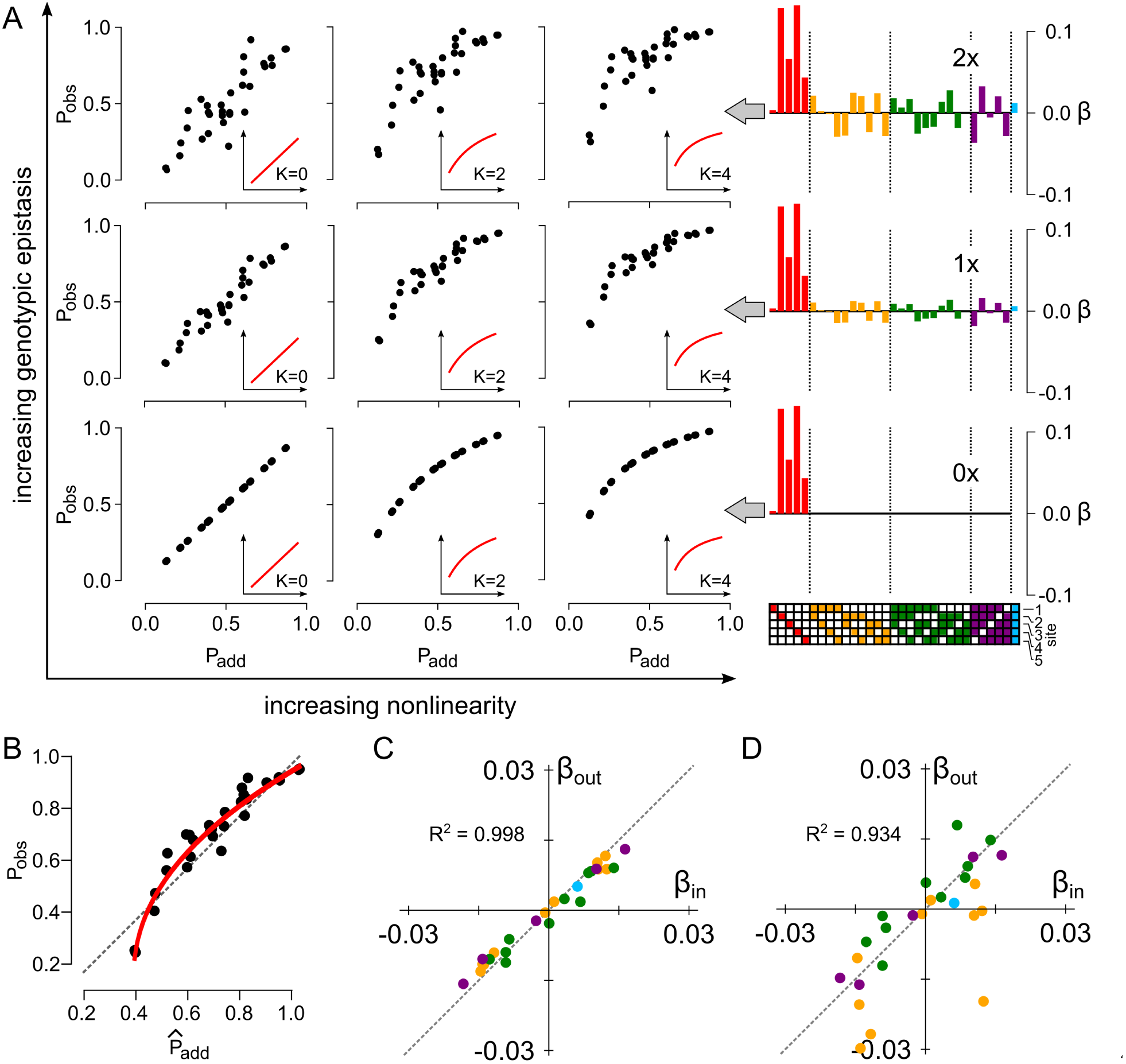
Epistasis and nonlinear scale induce different patterns of nonadditivity. A) Patterns of nonadditivity for increasing epistasis and nonlinear scale. Main panel shows a grid ranging from no epistasis, linear scale (bottom-left) to high epistasis, highly nonlinear scale (top-right). Insets in sub-panels show added nonlinearity. Going from left to right: *K* = 0, *K* = 2, *K* = 4. Epistatic coefficient plots to right show the magnitude of the input high-order epistasis, with colors and annotation as in Fig 2C. B) Plot of *P*_*obs*_ against 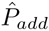 for the middle sub panel in panel A. Red line is the fit of the power transform to these data. C) Correlation between epistatic coefficients input into the simulation and extracted from the simulation after linearization by the power transform. Each point is an epistatic coefficient, colored by order. The Pearson’s correlation coefficient is shown in the upper-left quadrant. D) Correlation between epistatic coefficients input into the simulation and extracted from the simulation without application of the power transform.

### Nonlinearity can be separated from underlying high-order epistasis

The *P*_*obs*_ vs. *P*_*add*_ plots suggest an approach to disentangle high-order epistasis from nonlinear scale. By fitting a function to the *P*_*obs*_ vs *P*_*add*_ curve, we describe a transformation that relates the linear *P*_*add*_ scale to the (possibly nonlinear) *P*_*obs*_ scale (Schenk et al. 2013; Szendro et al. 2013). Once the form of the nonlinearity is known, we can then linearize the phenotypes so they are on an appropriate scale for epistatic analysis. Variation that remains (i.e. scatter) can then be confidently partitioned into epistatic coefficients.

In the absence of knowledge about the source of the nonlinearity, a natural choice is a power transform (Box and Cox 1964; Carroll and Ruppert 1981), which identifies a monotonic, continuous function through *P*_*obs*_ vs. *P*_*add*_. A key feature of this approach is that power-transformed data are normally distributed around the fit curve and thus appropriately scaled for regression of a linear epistasis model.

We tested this approach using one of our simulated data sets. One complication is that, for an experimental map, we do not know Padd. In the analysis above, we determined Padd from the additive coefficients used to generate the space. In a real map, Padd is not known; therefore, we had to estimate Padd. We did so by measuring the average effect of each mutation across all backgrounds, and then calculating 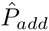 for each genotype as the sum of these average effects (Eq.1)

We fit the power transform to *P*_*obs*_ vs. 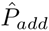 (solid red line, Fig 3B). The curve captures the nonlinearity added in the simulation. We linearized *P*_*obs*_ using the fit model (Eq. 2), and then extracted high-order epistatic coefficients. The extracted coefficients were highly correlated with the coefficients used to generate the map (*R*^2^ = 0:998) (Fig 3C). In contrast, applying the linear epistasis model to this map without first accounting for nonlinearity gives much greater scatter between the input and output coefficients (*R*^2^ = 0:934) (Fig 3D). This occurs because phenotypic variation from nonlinearity is incorrectly partitioned into the linear epistatic coefficients.

### Nonlinearity is a common feature of genotype-phenotype maps

Our next question was whether experimental maps exhibited nonlinear scales. We selected seven genotype-phenotype maps that had previously been reported to exhibit high-order epistasis (Table 1) and fit power transforms to each dataset (Fig 4, S1). We expected some phenotypes to be multiplicative (e.g. datasets I, II and IV were relative fitness), while we expected some to be additive (e.g. dataset IV is a free energy). Rather than rescaling the multiplicative datasets by taking logarithms of the phenotypes, we allowed our power transform to capture the appropriate scale. The power-transform identified nonlinearity in the majority of data sets. Of the seven data sets, three were less-than-additive (II, V, VI), two were greater-than-additive (III, IV), and two were approximately linear (I, VII). All data sets gave random residuals after fitting the power transform (Fig 4, S1).

**Fig 4:**
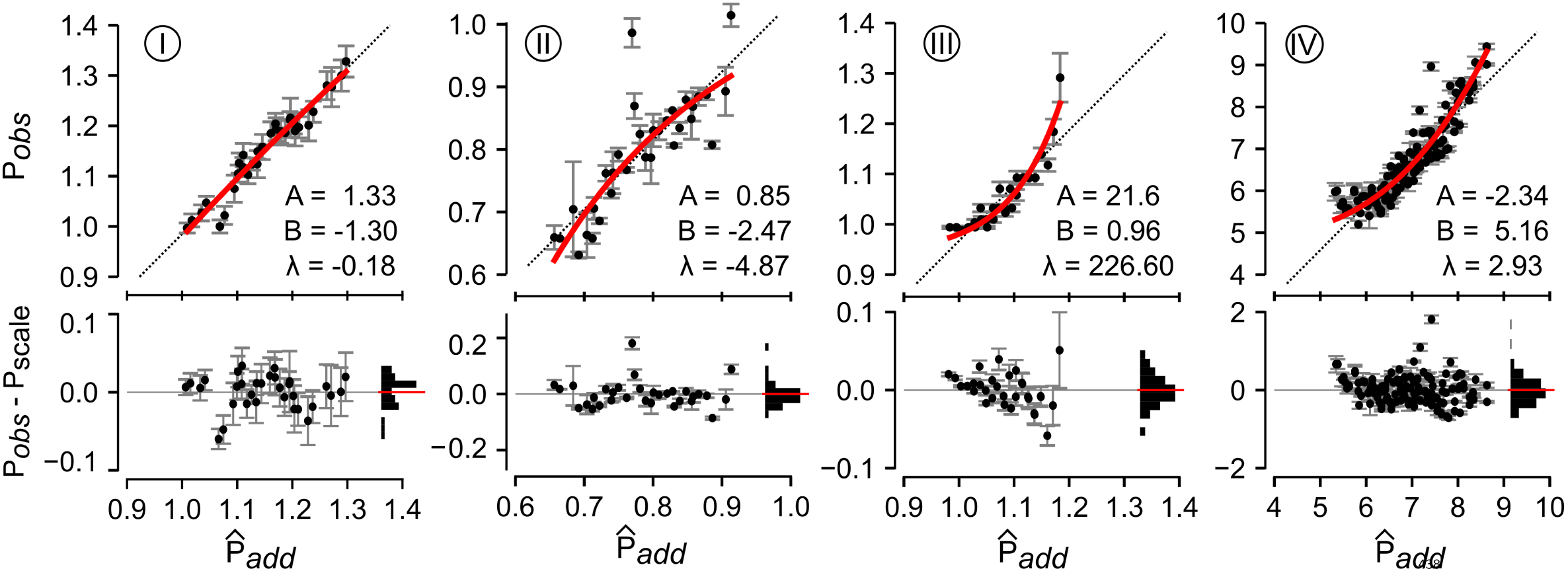
Experimental genotype-phenotype maps exhibit nonlinear phenotypes. Plots show observed phenotype *P*_*obs*_ plotted against 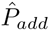 (Eq. 1) for data sets I through IV. Points are individual genotypes. Error bars are experimental standard deviations in phenotype. Red lines are the fit of the power transform to the data set. Pearson’s coefficient for each fit are shown on each plot. Dashed lines are *P*_*add*_ = *P*_*obs*_. Bottom panels in each plot show residuals between the observed phenotypes and the red fit line. Points are the individual residuals. Errorbars are the experimental standard deviation of the phenotype. The horizontal histograms show the distribution of residuals across 10 bins. The red lines are the mean of the residuals.

### High-order epistasis is a common feature of genotype-phenotype maps

With estimated scales in hand, we linearized the maps using Eq. 2 and re-measured epistasis (Fig S2). We used bootstrap sampling of uncertainty in the measured phenotypes to determine the uncertainty of each epistatic coefficient (see Methods), and then integrated these distributions to determine whether each coefficient was significantly different than zero. We then applied a Bonferroni correction to each p-value to account for multiple testing.

Despite our conservative statistical approach, we found high-order epistasis in every map studied (Fig 5A, S3). Every data set exhibited at least one statistically significant epistatic coefficient of fourth order or higher. We even detected statistically significant fifth-order epistasis (blue bar in Fig 5A, data set II). High-order coefficients were both positive and negative, often with magnitudes equal to or greater than the second-order terms. These results reveal that high-order epistasis is a robust feature of these maps, even when nonlinearity and measurement uncertainty in the genotype-phenotype map is taken into account.

We also dissected the relative contributions of each epistatic order to the remaining variation. To do so, we created truncated epistasis models: an additive model, a model containing additive and pairwise terms, a model containing additive through third-order terms, etc. We then measured how well each model accounted for variation in the phenotype using a Pearson’s coefficient between the fit and the data. Finally, we asked how much the Pearson coefficient changed with addition of more epistatic coefficients. For example, to measure the contribution of pairwise epistasis, we took the difference in the correlation coefficient between the additive plus pairwise model and the purely additive model.

The contribution of epistasis to the maps was highly variable (Fig 5B, S3). For data set I, epistatic terms explained 5.9% of the variation in the data. The contributions of epistatic coefficients decayed with increasing order, with fifth-order epistasis only explaining 0.1% of the variation in the data. In contrast, for data set II, epistasis explains 43.3% of the variation in the map. Fifth-order epistasis accounts for 6.3% of the variation in the map. The other data sets had epistatic contributions somewhere between these extremes.

### Accounting for nonlinear genotype-phenotype maps alters epistatic coefficients

Finally, we probed to what extent accounting for nonlinearity in phenotype altered the epistatic coefficients extracted from each space. Fig 6 and S4 show correlation plots between epistatic coefficients extracted both with and without linearization. The first-order coefficients were all highly correlated between the linear and nonlinear analyses for all data sets (Fig S5).

**Fig 5:**
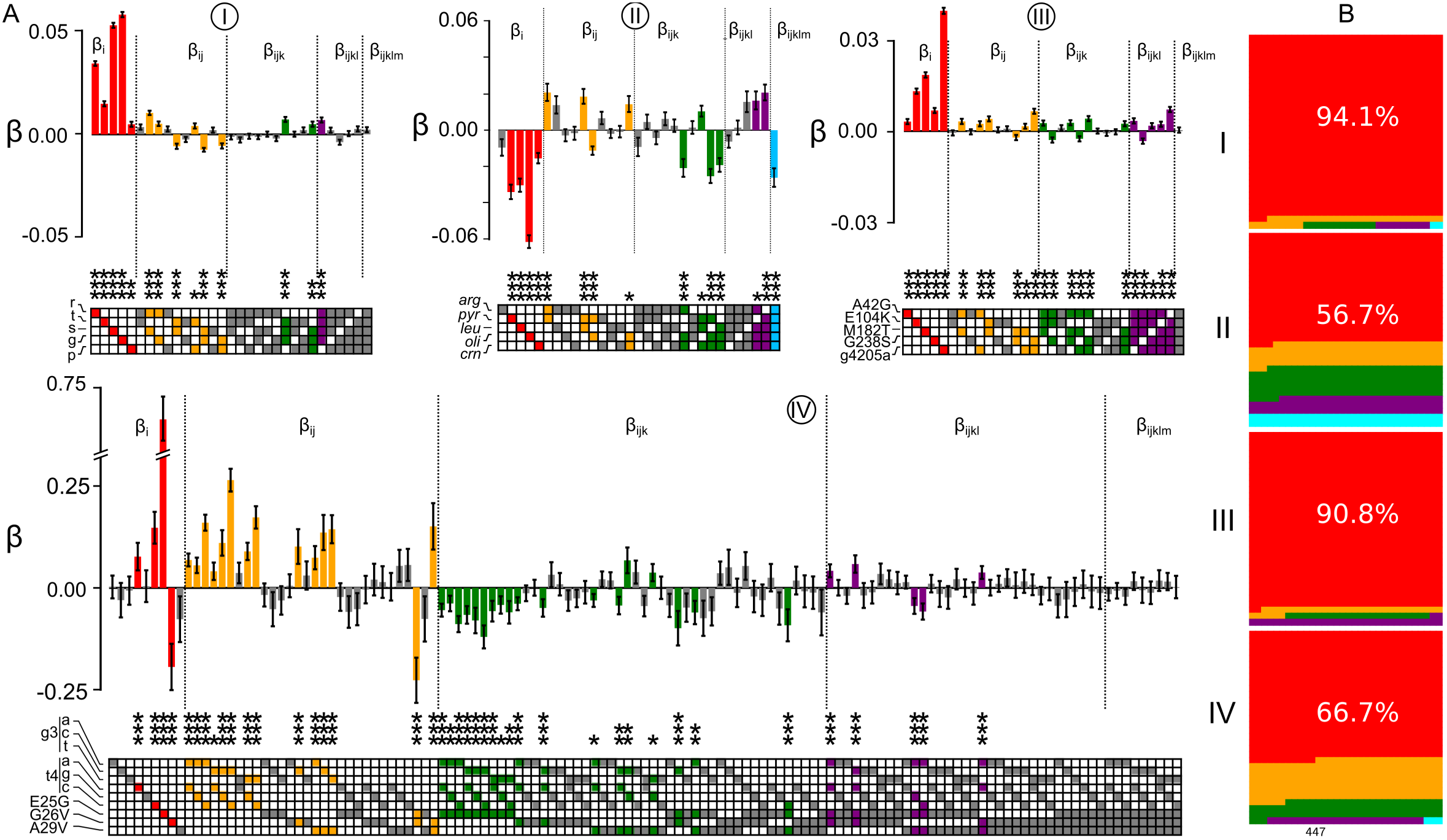
High-order epistasis is present in genotype-phenotype maps. A) Panels show epistatic coefficients extracted from data sets I-IV (Table 1, data set label circled above each graph). Bars denote coefficient magnitude and sign; error bars are propagated measurement uncertainty. Color denotes the order of the coefficient: first (*β*_*i*_, red), second (*β*_*ij*_, orange), third (*β*_*ijk*_, green), fourth (*β*_*ijkl*_, purple), and fifth (*β*_*ijklm*_, blue). Bars are colored if the coefficient is significantly different than zero (Z-score with *p-*value < 0.05 after Bonferroni correction for multiple testing). Stars denote relative significance: *p <* 0.05 (*), *p <* 0.01 (**), *p <* 0.001 (***). Filled squares in the grid below the bars indicate the identity of mutations that contribute to the coefficient. The names of the mutations, taken from the original publications, are indicated to the left of the grid squares. B) Sub-panels show fraction of variation accounted for by first through fifth order epistatic coefficients for data sets I-IV (colors as in panel A). Fraction described by each order is proportional to area.

**Fig 6:**
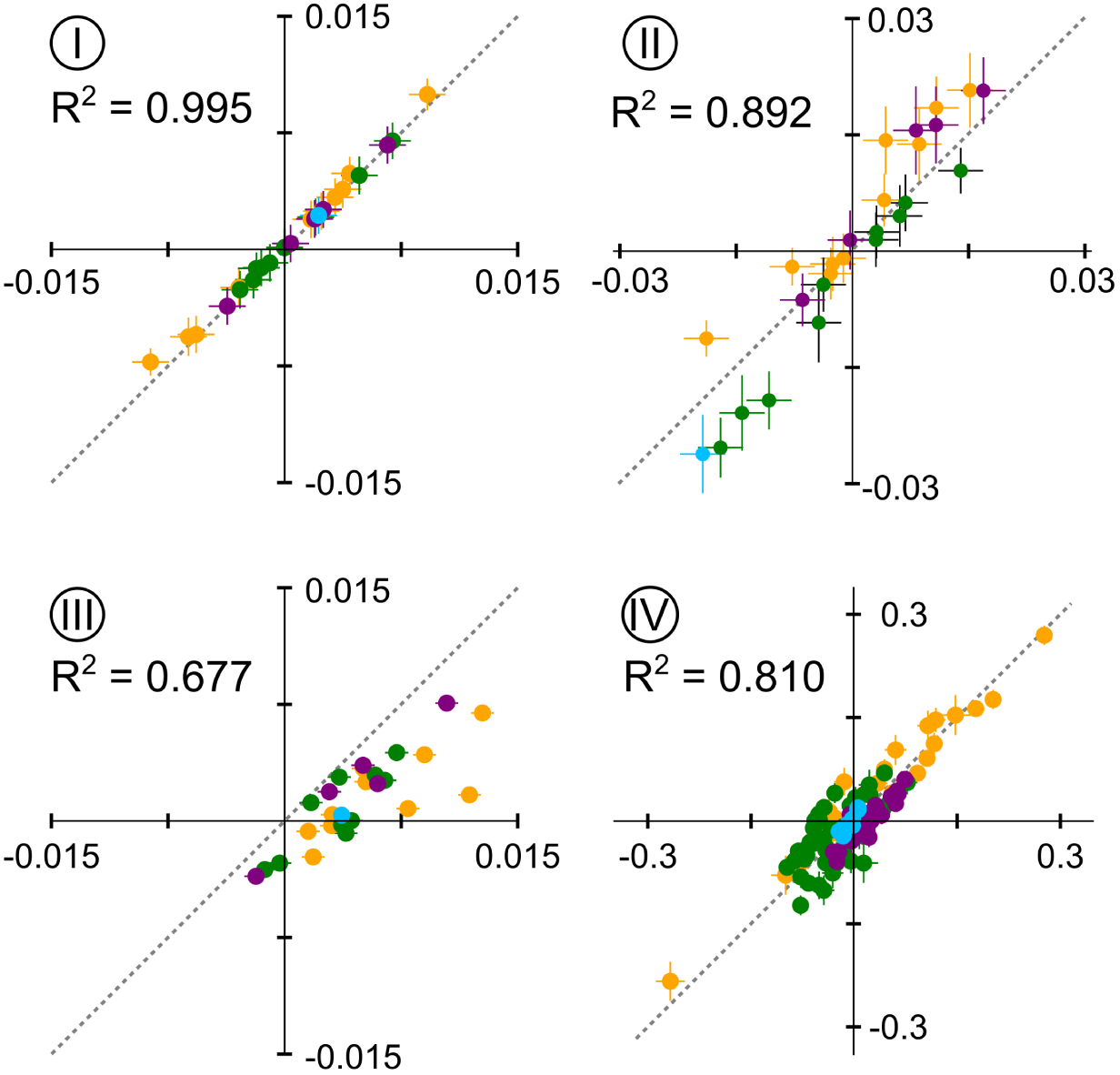
Nonlinear phenotypes distort measured epistatic coefficients. Sub-panels show correlation plots between epistatic coefficients extracted without accounting for nonlinearity (*x*-axis) and accounting for linearity (*y*-axis) for data sets I-IV. Each point is an epistatic coefficient, colored by order. Error bars are standard deviations from bootstrap replicates of each fitting approach.

For the epistatic coefficients, the degree of correlation depended on the degree of nonlinearity in the dataset. Data set I—which was essentially linear—had identical epistatic coefficients whether the nonlinear scale was taken into account or not. In contrast, the other data sets exhibited scatter off of the line. Data set III was particularly noteworthy. The epistatic coefficients were systematically overestimated when the nonlinear scale was ignored. Two large and favorable pairwise epistatic terms in the linear analysis became essentially zero when nonlinearity was taken into account. These interactions—M182T/g4205a and G283S/g4205a—were both noted as determinants of evolutionary trajectories in the original publication (Weinreich et al. 2006); however, our results suggest the interaction is an artifact of applying a linear model to a nonlinear data set. Further ≈ 20% (six of 27) epistatic coefficients flipped sign when nonlinearity was taken into account (Fig 6, III, bottom right quadrant).

Overall, we found that low-order epistatic coefficients were more robust to the linear assumption than high-order coefficients. Data set IV is a clear example of this behavior. The map exhibited noticeable nonlinearity (Fig 4). The first-and second-order terms were well correlated between the linear and nonlinear analyses (Fig 6, S4, S5). Higher-order terms, however, exhibited much poorer overall correlation. While the R^2^ for second-order coefficients was 0.95, the correlation was only 0.43 for third-order. This suggests that previous analyses of nonlinear genotype-phenotype maps correctly identified the key mutations responsible for variation in the map, but incorrectly estimated the high-order epistatic effects.

## Discussion

Our results reveal that both nonlinear scales and high-order epistasis play important roles in shaping experimental genotype-phenotype maps. Five of the seven data sets we investigated exhibited nonlinear scales, and all of the data sets exhibited high-order epistasis, even after accounting for non-linearity. This suggests that both should be taken into account in analyses of genotype-phenotype maps.

### Origins of nonlinear scales

We observed two basic forms of nonlinearity in these maps: saturating, less-than-additive maps and exploding, greater-than-additive maps. Many have observed less-than-additive maps in which mutations have lower effects when introduced into more optimal backgrounds (MacLean et al. 2010; Chou et al. 2011). Such saturation has been proposed to be a key factor shaping evolutionary trajectories (MacLean et al. 2010; Chou et al. 2011; Kryazhimskiy et al. 2014; Tokuriki et al. 2012; Otto and Feldman 1997). Further, it is intuitive that optimizing a phenotype becomes more difficult as that phenotype improves. Our nonlinear fits revealed this behavior in three different maps.

The greater-than-additive maps, in contrast, were more surprising: why would mutations have a larger effect when introduced into a more favorable background? For the *β*-lactamase genotype-phenotype map (III, Fig 4), it appears this is an artifact of the original analysis used to generate the data set (Weinreich et al. 2006). This data set describes the fitness of bacteria expressing variants of an enzyme with activity against *β*-lactam antibiotics. The original authors measured the minimum-inhibitory concentration (MIC) of the antibiotic against bacteria expressing each enzyme variant. They then converted their MIC values into apparent fitness by sampling from an exponential distribution of fitness values and assigning these fitness values to rank-ordered MIC values (Weinreich et al. 2006). Our epistasis model extracts this original exponential distribution (Fig S6). This result demonstrates the effectiveness of our approach in extracting nonlinearity in the genotype-phenotype map.

The origins of the growth in the transcription factor/DNA binding data set are less clear (IV, Fig 4). The data set measures the binding free energy of variants of a transcription factor binding to different DNA response elements. We are aware of no physical reason for mutations to have a larger effect on free energy when introduced into a background with better binding. One possibility is that the genotype-phenotype map reflects multiple features that are simultaneously altered by mutations, giving rise to this nonlinear shape. This is a distinct possibility in this data set, where mutations are known to alter both DNA binding affinity and DNA binding cooperativity (McKeown et al. 2014).

### Best Practice

Because nonlinearity is a common feature of these maps, linearity should not be assumed in analyses of epistasis. Given a sufficient number of phenotypic observations, however, the appropriate scale can be estimated by construction of a *P*_*obs*_ vs. *P*_*add*_ plot and regression of a nonlinear scale model. With this scale in hand, one can then transform the genotype-phenotype map onto a linear scale appropriate for analysis using a high-order epistasis model. Our software pipeline automates this process. It takes any genotype-phenotype map in a standard text format, fits for nonlinearity, and then estimates high-order epistasis. It is freely available for download (https://harmslab.github.com/epistasis).

One important question is how to select an appropriate function to describe the nonlinear scale. By visual inspection, all of the data sets we studied were monotonic in 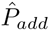 and could be readily captured by a power transform. Other maps may be better captured with other functions. For example, inspection of a *P*_*obs*_ vs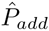 plot could reveal a non-monotonic scale, leading to a better fit with a polynomial than a power-transform. Another possibility is that external biological knowledge motivates scale choice (Schenk et al. 2013).

The choice of model determines what fraction of the variation is assigned to “scale” versus “epistasis.” The more complicated the function chosen, the more variation in the data is shifted from epistasis and into scale. One could, for example, fit a completely uninformative *Lth*-order polynomial, which would capture all of variation as scale and none as epistasis. Scale estimation should governed by the well-established principles of model regression: find the simplest function that captures the maximum amount of variation in the data set without fitting stochastic noise. Because epistasis is scatter off the scale line (noise), model-selection approaches like the F-test, Akaike Information Criterion, and inspection of fit residuals are a natural strategy for partitioning variation between scale and epistasis.

### Interpretation

Another powerful aspect of this approach is that it allows explicit separation of two distinct origins of non-additivity in genotype-phenotype maps.

This can be illustrated with a simple, conceptual, example. Imagine mutations to an enzyme, expressed in bacteria, that have a less-than-additive effect on bacterial growth rate. To a first approximation, this epistasis could have two origins. The first is at the level of the enzyme: maybe the mutations have a specific, negative chemical interactions that alter enzyme rate. The second is at the level of the whole cell: maybe, above a certain activity, the enzyme is fast enough that some other part of the cell starts limiting growth. Mutations continue to improve enzyme activity, but growth rate does not reflect this. These two origins of less-than-additive behavior will have different effects in a *P*_*add*_ vs. *P*_*obs*_ plot: saturation of growth rate will appear as nonlinearity, interactions between mutations at the enzyme level will appear as linear epistasis. Our analysis would reveal this pattern and set up further experiments to tease apart these possibilities.

This may also provide important evolutionary insights. One important question is to what extent evolutionary paths are shaped by global constraints versus specific interactions that lead to specific historical contingencies (Harms and Thornton 2014; Shah et al. 2015; Kryazhimskiy et al. 2014). For example, recent work has shown specific epistatic interactions lead to sequence-level unpredictability, while a globally less-than-additive scale leads to predictable phenotypes in evolution (Kryazhimskiy et al. 2014). Our analysis approach naturally distinguishes these origins of non-additivity, and thus these evolutionary possibilities. Prevailing magnitude epistasis (de Visser et al. 2009), global epistasis (Kryazhimskiy et al. 2014), and diminishing-returns epistasis (Chou et al. 2011; MacLean et al. 2010; Otto and Feldman 1997; Tokuriki et al. 2012) will all appear as nonlinear scales. In contrast, specific interactions will appear in specific coefficients in the linear epistasis model. Our detection of nonlinearity and high-order epistasis in most datasets suggests that both forms of non-additivity will be in play over evolutionary time.

### High-order epistasis

Finally, our work reveals that high-order epistasis is, indeed, a common feature of genotypephenotype maps. Our study could be viewed as an attempt to “explain away” previously observed high-order epistasis. To do so, we both accounted for nonlinearity in the map and propagated experimental uncertainty to the epistatic coefficients. Surprisingly—to the authors, at least—high-order epistasis was robust to these corrections.

High-order epistasis can make huge contributions to genotype-phenotype maps. In data set II, third-order and higher epistasis accounts for fully 31.0% of the variation in the map. The average contribution, across maps, is 12.7%. We also do not see a consistent decay in the contribution of epistasis with increasing order. In data sets II, V and VI, third-order epistasis contributes more variation to the map than second-order epistasis. This suggests that epistasis could go to even higher orders in larger genotype-phenotype maps.

The generality of these results across all genotype-phenotype maps is unclear. The maps we analyzed were measured and published because they were “interesting,” either from a mechanistic or evolutionary perspective. Further, most of the maps have a single, maximum phenotype peak. The nonlinearity and high-order epistasis we observed may be common for collections of mutations that, together, optimize a function, but less common in “flatter” or more random genotype-phenotype maps. This can only be determined by characterization of genotype-phenotype maps with different structural features.

The observation of this epistasis also raises important questions: What are the origins of third, fourth, and even fifth-order correlations in these data sets? What, mechanistically, leads to a five-way interaction between mutations? Does neglecting high-order epistasis bias estimates of low-order epistasis (Otwinowski and Plotkin 2014)? What can this epistasis tell us about the biological underpinning of these maps (Lehár et al. 2008; Hu et al. 2011, 2013; Taylor and Ehrenreich 2015)?

The evolutionary implications are also potentially fascinating. Epistasis creates temporal dependency between mutations: the effect of a mutation depends strongly on specific mutations that fixed earlier in time (Bedau and Packard 2003; Desai 2009; Harms and Thornton 2014; Shah et al. 2015) How does this play out for high-order epistasis, which introduces long-range correlations across genotype-phenotype maps? Do these low magnitude interactions matter for evolutionary outcomes or dynamics? These, and questions like them, are challenging and fascinating future avenues for further research.

## Acknowledgments

We would like to thank Patrick Phillips, Jamie Bridgham and members of the Harms lab for helpful discussions and comments. We would also like to thank David Hall for providing the complete data for data sets VI and VII. Work was supported by start up funds from the University of Oregon (ZRS). MJH is a Pew Scholar in the Biomedical Sciences, supported by The Pew Charitable Trusts.

---

**Fig S1:**
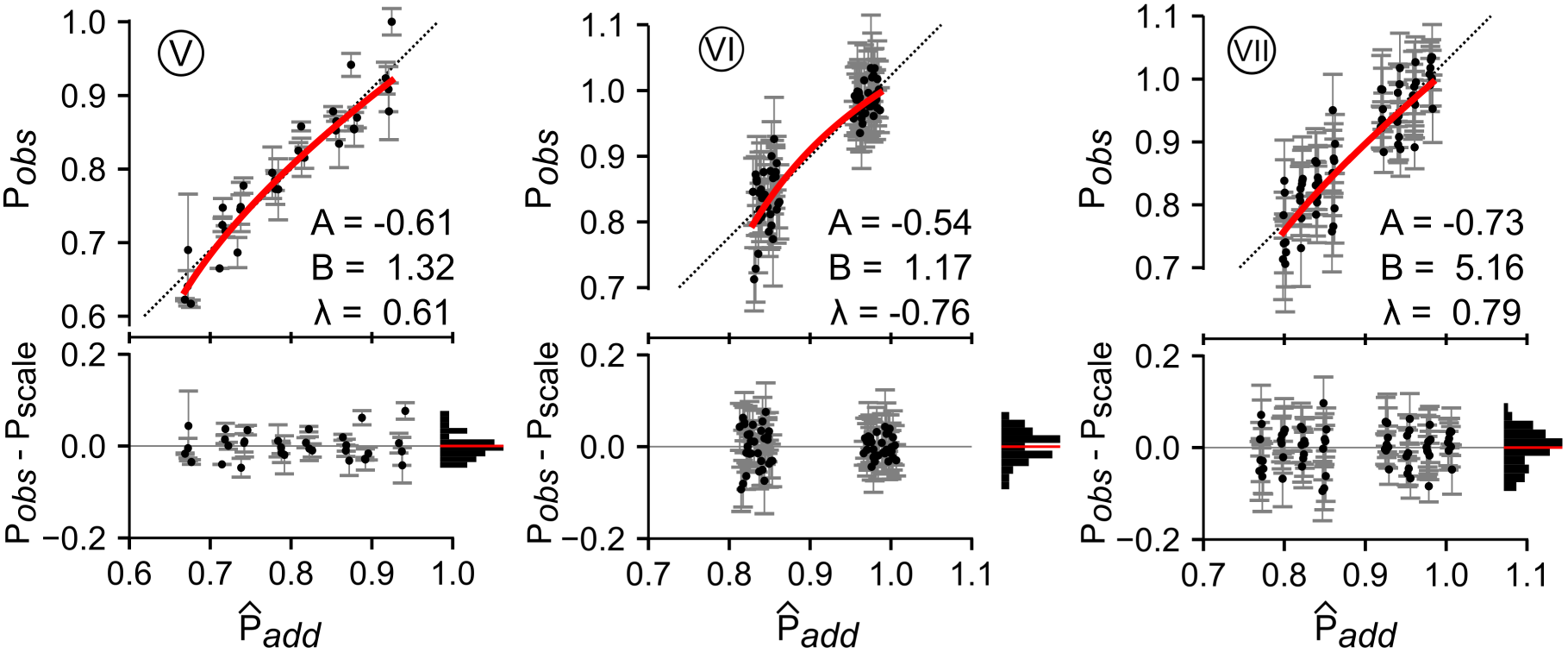
Experimental genotype-phenotype maps exhibit nonlinear phenotypes. Plots show observed phenotype *P*_*obs*_ plotted against 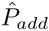 (Eq. 1) for data sets V through VII. Points are individual genotypes. Error bars are experimental standard deviations in phenotype. Red lines are the fit of the power transform to the data set. Pearson’s coefficient for each fit are shown on each plot. Dashed lines are *P*_*add*_ = *P*_*obs*_. Bottom panels in each plot show residuals between the observed phenotypes and the red fit line. Points are the individual residuals. Errorbars are the experimental standard deviation of the phenotype. The horizontal histograms show the distribution of residuals across 10 bins. The red lines are the mean of the residuals.

**Fig S2:**
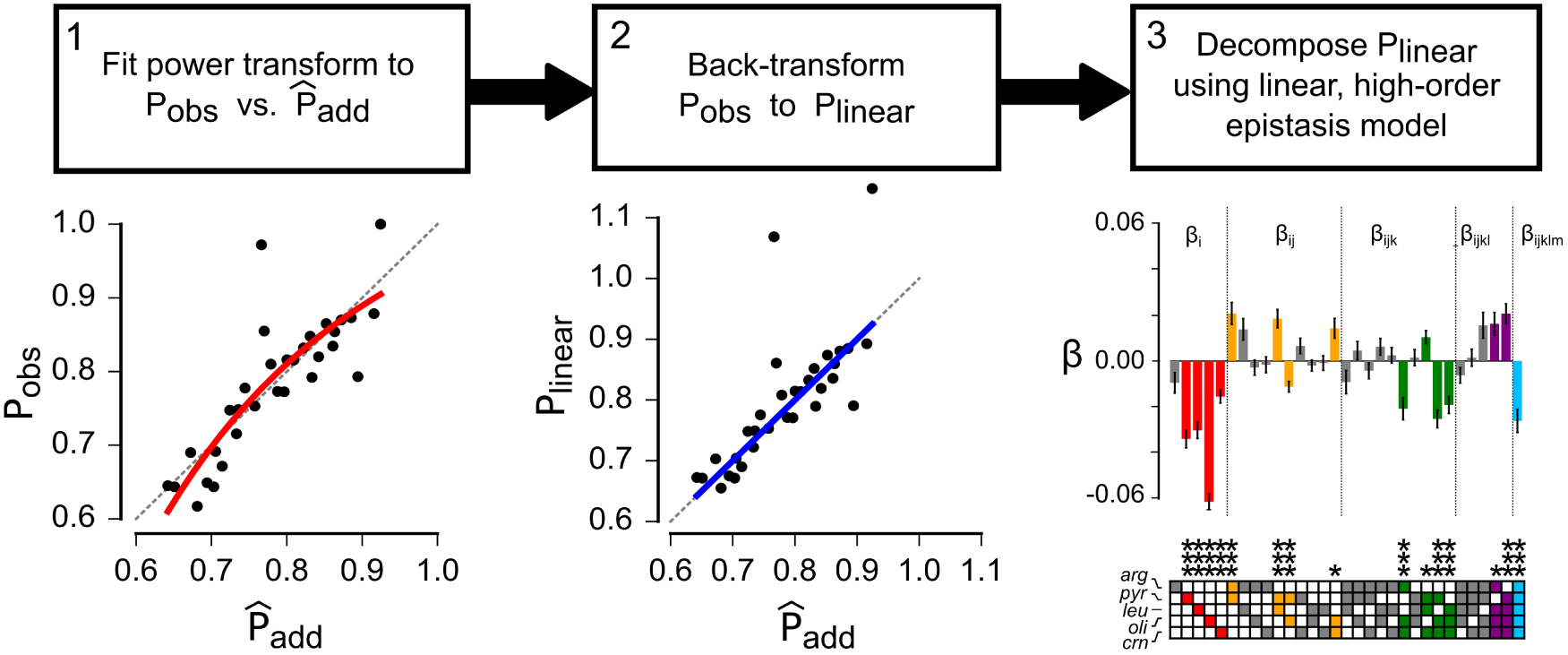
Nonlinear phenotypes can be transformed to linear scale to estimate high-order epistasis. Flowchart shows the steps for estimating high-order epistasis in nonlinear genotype-phenotype maps. The plots beneath the chart show this pipeline for data set II. In step 1, a power transform function is used to fit the *P*_*obs*_ versus 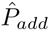 plot and estimate the map’s scale. In step 2, the inverse of the fitted transform is used to back-transform *P*_*o*_*b*_*s*_ to a linear scale, *P*_*linear*_. In step 3, a linear, high-order epistasis model is used to fit the variation in *P*_li_*_near_*. In the left plot, points are individual genotypes, red line is the resulting fit and dashed line is the *P*_*add*_ = *P*_*obs*_. In the middle plot, the blue line is the new scale of *P*_*obs*_ after back transforming. In the right plot, bars represent additive and epistatic coefficients extracted from the linear phenotypes. Error bars are propagated measurement uncertainty. Color denotes the order of the coefficient: first (*β*_*i*_, red), second (*β*_*ij*_, orange), third (*β*_*ijk*_, green), fourth (*β*_*ijkl*_, purple), and fifth (*β*_*ijklm*_, blue). Bars are colored if the coefficient is significantly different than zero (Z-score with p-value <0.05 after Bonferroni correction for multiple testing). Stars denote relative significance: p < 0.05 (*), p < 0.01 (**), p < 0.001 (***). Filled squares in the grid below the bars indicate the identity of mutations that contribute to the coefficient.

**Fig S3:**
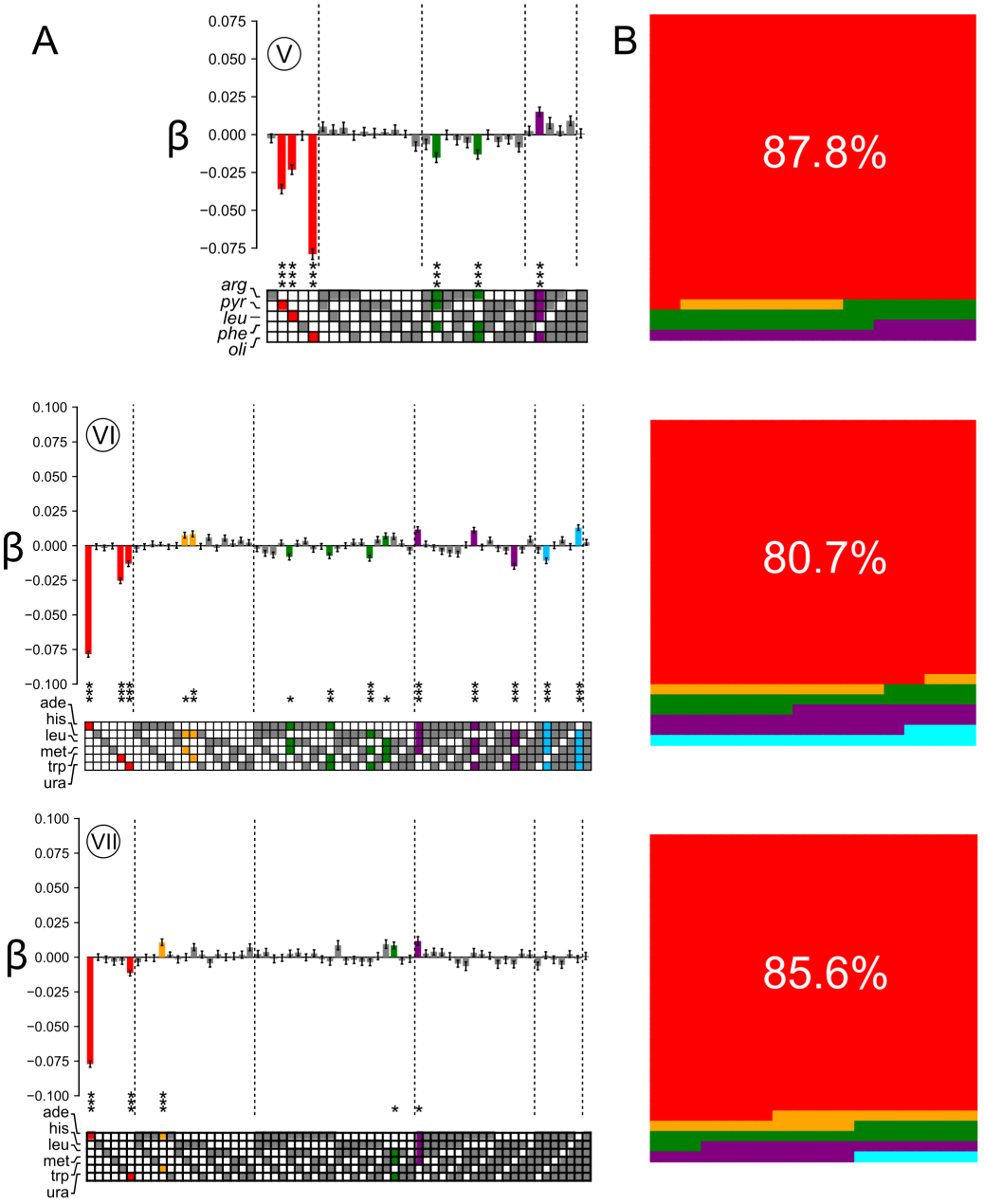
High-order epistasis is present in genotype-phenotype maps. A) Panels show epistatic coefficients extracted from data sets V-VII (Table 1, data set label circled above each graph). Bars denote coefficient magnitude and sign; error bars are propagated measurement uncertainty. Color denotes the order of the coefficient: first (*β*_*i*_, red), second (*β*_*ij*_, orange), third (*β*_*ijk*_, green), fourth (*β*_*ijkl*_, purple), and fifth (*β*_*ijklm*_, blue). Bars are colored if the coefficient is significantly different than zero (Z-score with p-value <0.05 after Bonferroni correction for multiple testing). Stars denote relative significance: p < 0.05 (*), p < 0.01 (**), p < 0.001 (***). Filled squares in the grid below the bars indicate the identity of mutations that contribute to the coefficient. The names of the mutations, taken from the original publications, are indicated to the left of the grid squares. B) Sub-panels show fraction of variation accounted for by first through fifth order epistatic coefficients for data sets I-IV (colors as in panel A). Fraction described by each order is proportional to area.

**Fig S4:**
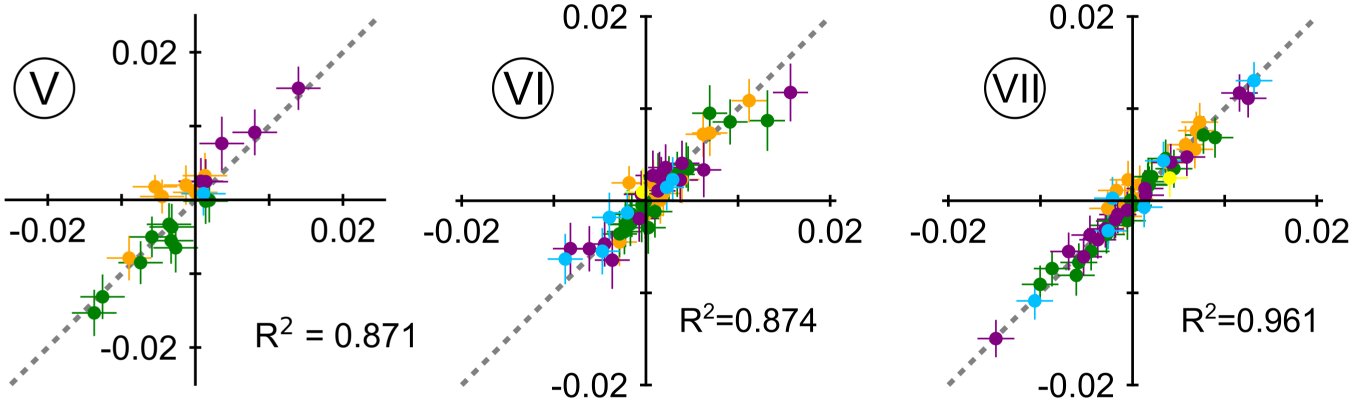
Nonlinear phenotypes distort measured epistatic coefficients. Sub-panels show correlation plots between epistatic coefficients extracted without accounting for nonlinearity (*x*-axis) and accounting for linearity (*y* -axis) for data sets V-VII. Each point is an epistatic coefficient, colored by order.

**Fig S5:**
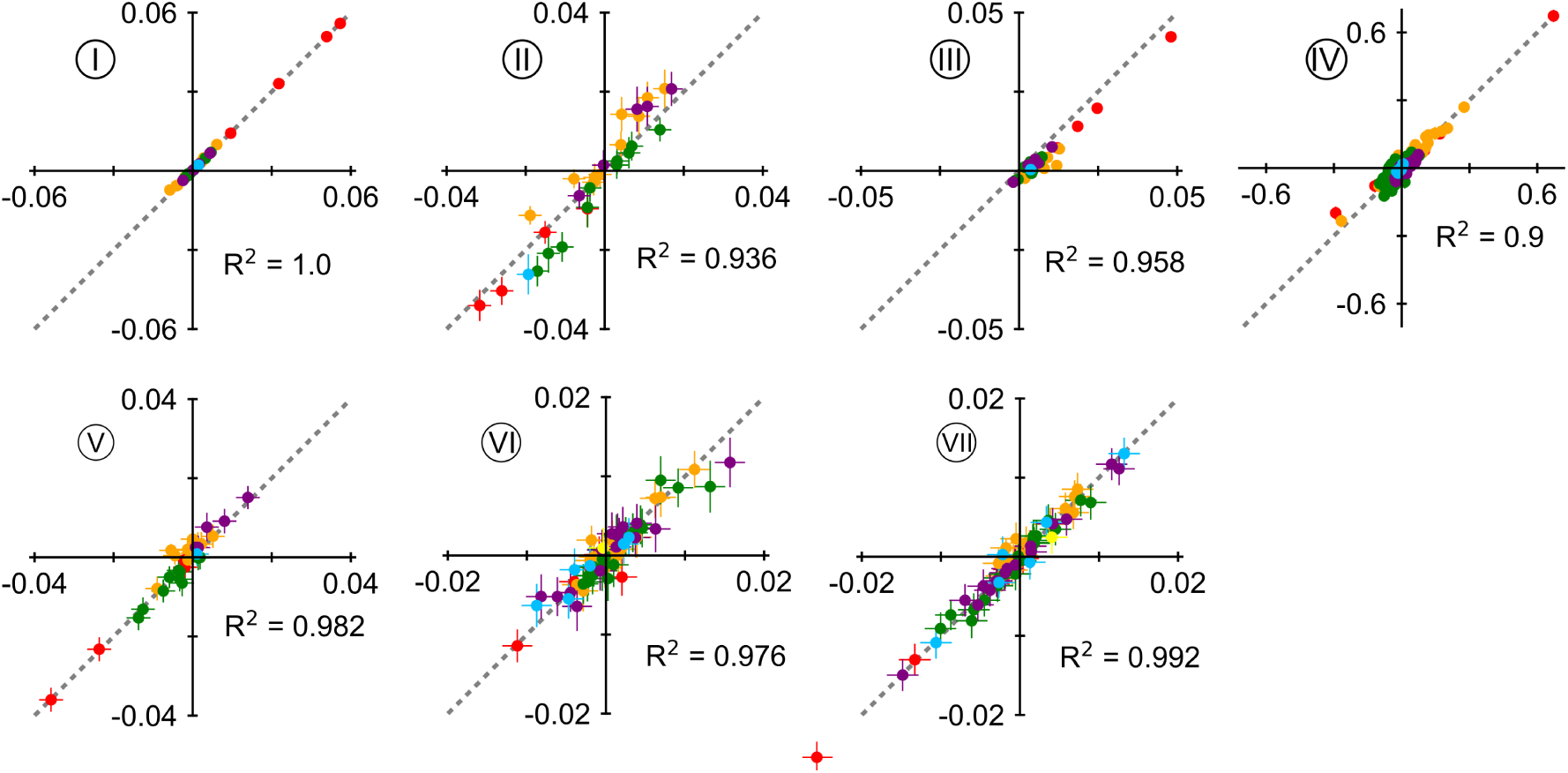
Additive coefficients are well estimated, even when nonlinearity is neglected. Sub-panels show correlation plots between both additive and epistatic coefficients extracted without accounting for nonlinearity (*x*-axis) and accounting for linearity (*y*-axis) for data sets I-VII. Each point is an epistatic coefficient, colored by order. Error bars are standard deviations from bootstrap replicates of each fitting approach.

**Fig S6:**
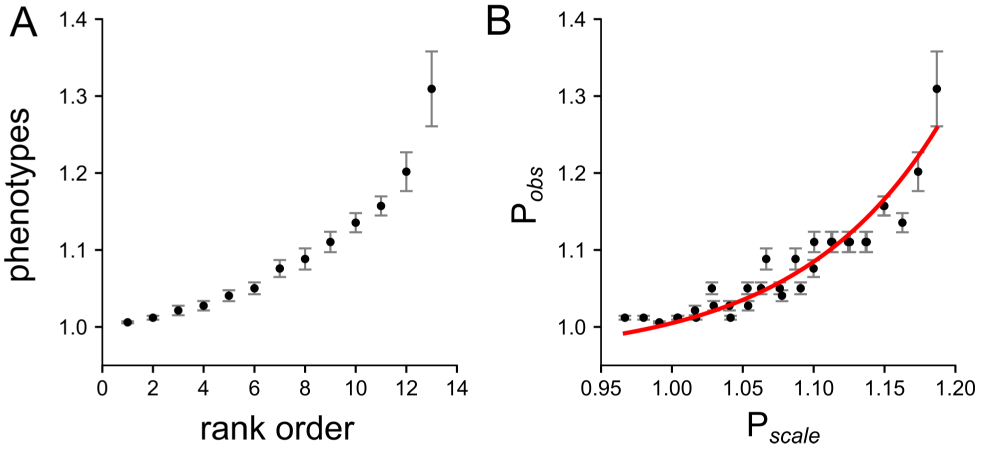
Exponential fitness model leads to global nonlinearity in the *β*-lactamase data set (III). A) A recapitulation of the map used in the original publication (Weinreich et al. 2006). We first rank-ordered the genotypes according to the measured property (the minimum inhibitory concentration of a *β*-lactam antibiotic against a clonal population of bacteria expressing that protein). This gave us 13 classes of genotypes, as some genotypes had equivalent MIC values. We then drew 3,000 random fitness values from the distribution *W* = 1+x, where *x* is an exponential distribution centered around 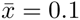. We took the top 13 values from this distribution and assigned them, in value order, to each of the 32 *β*-lactamase genotypes. Panel A shows the average and standard deviation of the fitness values *W* assigned to each of these ranks if we repeat the protocol above 1,000 times. B) Best fit for the power-transform for data set III. Solid red line denotes the best fit (nonlinear). This fit successfully pulls out the original distribution of *W.*

